# Secure federated Boolean count queries using fully-homomorphic cryptography

**DOI:** 10.1101/2021.11.10.468090

**Authors:** Alexander T. Leighton, Yun William Yu

**Affiliations:** Harvard Medical School, Boston MA 02115, USA; Brandeis University, Waltham MA 02453, USA; University of Toronto Scarborough, Toronto ON M1C 1A4, Canada; Carnegie Mellon University, Pittsburgh PA 15213, USA

**Keywords:** Federated count-query, Probabilistic sketch, Homomorphic encryption

## Abstract

Biomedical data is often distributed between a network of custodians, causing challenges for researchers wishing to securely compute aggregate statistics on those data without centralizing everything—the prototypical ‘count query’ asks how many patients match some multifaceted set of conditions across a network of hospitals. Difficulty arises from two sources: (1) the need to deduplicate patients who may be present in the records of multiple hospitals and (2) the need to unify partial records for the same patient which may be split across hospitals. Although cryptographic tools for secure computation promise to enable collaborative studies with formal privacy guarantees, existing approaches either are computationally impractical or support only simplified analysis pipelines. To the best of our knowledge, no existing practical secure method addresses both of these difficulties simultaneously.

Here, we introduce secure federated Boolean count queries using a novel 2-stage probabilistic sketching and sampling protocol that can be efficiently implemented in off-the-shelf federated homomorphic encryption libraries (Palisade and Lattigo), provably ensuring data security. To this end, we needed several key technological innovations, including re-encoding the LogLog union-cardinality sketch and designing an appropriate sampling for intersection cardinalities. Our benchmarking shows that we can answer federated Boolean count queries in less than 2 CPU-minutes with absolute errors in the range of 6% of the total number of touched records, while revealing only the final answer and the total number of touched records. With modern core-parallelism, we can thus answer queries on the order of seconds. Our study demonstrates that by computing on compressed and encrypted data, it is possible to securely answer federated Boolean count queries in real-time.

## 1 Introduction

The federated Boolean count query asks how many unique patients in a network match a given set of criteria [20]. Computing a cardinality is algorithmically straight-forward; however, as patient information is confidential, ideally statistics should be computed privately, protected even from other hospitals in the network. Two major problems arise when trying to compute aggregate statistics without revealing information about individuals: (1) it is difficult to resolve duplicate patients across institutions, and (2) it is difficult to combine partial information about the same patient across institutions. To illustrate, suppose we are interested in the question of how many patients at Beth Israel Deaconess Medical Center (BIDMC) and at Massachusetts General Hospital (MGH) have both hypertension and diabetes. If a patient Alice has records for hypertension and diabetes at both BIDMC and MGH, she may be double-counted. On the other hand, a patient Bob who is recorded at BIDMC only for hypertension and at MGH only for diabetes may not be counted at all, unless those records can be combined.

Note that because any Boolean query can be written in conjunctive normal form (an “AND of ORs”), we can rewrite any Boolean count as an intersection of unions. Thus, more precisely, given *n* parties and *n*_*c*_ conditions, let *A*_*i,j*_ be the records at party *i* matching condition *j*. Then, it is sufficient to be able to compute

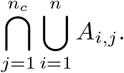

For example, if we want to know all patients who broke their legs and either have hypertension or diabetes (or both), we can rewrite that query as the union of hypertension and diabetes patients, intersected with the set of patients who broke their legs.

A growing body of work seeks to address this issue, but all practical approaches we have seen either rely on a trusted third party or reveal some private information. It is possible to control this data leakage with data use agreements, financial penalties, etc., but those are legal controls, not technical ones [18]. In prior work, we showed that probabilistic cardinality estimators (such as HyperLogLog [8]) can be used to trade off accuracy for privacy [21], as measured by the metric of k-anonymity [16]. Intuitively, probabilistic sketches and other lossy compression algorithms inherently add noise to the system, which may decrease the ability of an adversary to pinpoint an individual. Unfortunately, Desfontaines et al [5] proved that an adversary with access to all intermediate HyperLogLog probabilistic sketch representations during computation can reconstruct a large fraction of records. Furthermore, even a weaker adversary with only access to the partial sketches each hospital sends to a central party can reveal substantial amounts of information [17]. Stronger privacy guarantees can be had by combining sketching with differential privacy [7] on the intermediate sketches [14], but the noise-based mechanisms do still leak a quantifiable amount of private information, and furthermore decrease the accuracy of the final results.

Modern cryptographic tools can in theory resolve these shortcomings through fully homomorphic encryption [11]. Instead of adding noise to the data, all sensitive computations are run on ciphertext (encrypted data) and only the final answer is revealed; this is in many ways both simpler and more secure than the noise-based mechanisms when properly implemented—no data is leaked by intermediate computation. The typical Achilles heel of MPC is its speed—to compare, even the fast secure MPC GWAS protocols require hours of total single-threaded compute [2,9], perhaps reducible to tens of minutes using parallelization. While suitable for complex analyses, such runtimes exceed reasonable expectations in the workflow of clinical query systems, such as i2b2 or Shrine, which are expected to return queries on the order of only a few minutes at most. The recent MPC-FM [13] and DP-DICE [19] algorithms fully solve what they call the secure distributed cardinality estimation problem, the latter on the order of minutes, which is very nearly fast enough. However, the secure distributed cardinality estimation task only resolves duplicate records across data custodians, and does not allow for combining partial information about the same patient, so those do not solve the federated Boolean count query problem we pose—in the notation above, these methods only solve the union half of the problem, but not the intersection portion. Indeed, to our knowledge, there are no efficient protocols for the full federated Boolean count query problem.

In this work, we introduce secure federated Boolean count queries using a novel 2-stage probabilistic sketching and sampling protocol implemented using fully-homomorphic ring-learning with errors cryptography. In particular, we use the residue number system (RNS) variant of BFV encryption [12], as implemented in the Palisade [15] and Lattigo [1] software libraries. Our algorithm works by first computing an approximate count query of all “touched records” from encrypted LogLog [6] sketches; one key conceptual advance is in determining that unlike other sketches, a unary encoding of the LogLog sketches can be manipulated efficiently in ciphertext. This provides a very fast resolution to the secure distributed cardinality estimation problem (also solved by MPC-FM and DP-DICE). Then, we use the results of that count-query to determine the density at which to perform a blind coordinated random subsampling of the touched records at each hospital to estimate an intersection cardinality.

## 2 Methods

The maturity of fully homomorphic cryptographic libraries means that it is now feasible to treat them as black-box alternate computation platforms. A webdesigner need not understand the intricacies behind the HTTPS protocol, but can simply specify the security/computation tradeoff they want. A data scientist using GPU-acceleration needs only know that matrix multiplication and vectorized operations are faster, but conditionals are much slower. Similarly, with Palisade and Lattigo, algorithms designers can choose relevant parameters, and then are given a set of primitives around which to design algorithms, abstracting away the underlying cryptography.

### 2.1 Union cardinality

We begin by giving our implementation of secure union cardinality. Although existing work like MPC-FM [13] and DP-DICE [19] also solve this problem, that work was concurrent to ours, and ours is both easy to implement and fairly efficient.

Most work tackling privacy-preserving multiset cardinality estimation relies on the LogLog-family of cardinality estimators [6,8]. The basic idea behind these estimators is that given *n* uniformly random numbers between 0 and 1, the expected minimum of these numbers of 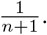 Thus, one naive way to estimate cardinality is to hash elements in an arbitrary set to uniformly random numbers, and then take the inverse of the minimum such value. Assuming we are consistent with our choice of hash function, we automatically get de-duplication because duplicate elements hash to the same value.

The LogLog family of algorithms reformulates the procedure to reduce both space-complexity and variance. First, instead of storing the minimum hash value exactly, one only needs to store the order-of-magnitude of the minimum hash values, which are likewise already logarithmic in the size of the set through binary encoding. Importantly, the double logarithmic space compression allows us to later use an exponential blow-up through a unary encoding of the bit vectors while still maintaining reasonable logarithmic space.

Another key idea from the literature is that keeping track of multiple minimum hash values for a set of buckets reduces variance; these buckets are purely an algorithmic feature unrelated to any clinical queries. In the LogLog algorithm, a fixed number of buckets, *m*, is picked. Then, elements are assigned a random hash value and bucket. We keep track of the minimum for each bucket, taking the combined minimum per bucket when merging into an aggregate approximation. Much of the rich historic work in the development of the LogLog family has aimed to construct lower-variance estimators. The original LogLog estimator uses an arithmetic mean of bucket values, and had a standard error of around 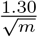. SuperLogLog improves upon LogLog by truncating extreme data, getting an effective error of around 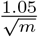. The most popular today, HyperLogLog, replaces the arithmetic mean of LogLog with a harmonic mean, with standard error 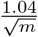. Unfortunately, neither conditional truncation nor harmonic means are efficient on the platform of modern homomorphic encryption, which really only efficiently supports addition and a few multiplications. As such, we realized we needed to switch to the older LogLog, which is less efficient on normal CPUs, but more efficient on ciphertext.

Our algorithm takes as input the query and each hospital returns *m* unary vectors representing a party’s sketches of the query. These sketches are encrypted using a multiparty public key, so that all of the hospitals have to coordinate in order to decrypt anything. Importantly, the hash values for LogLog are integers between 0 and 31, so we encode 0 as a vector of 32 ciphertext 1’s, 1 as a vector with a single 0 and then 31 1’s, and so on. If 5 is encoded as 0000011 …, and 2 is encoded as 0011111 … (counting the number of initial 0’s), then (5, 2) = 0000011 … × 0011111 = 0000011 …, which correctly encodes 5. This multiplication is the key to homomorphically merging LogLog sketches from different hospitals, as this operation lets us compute the maximum value in a particular bucket across sketches using log(*n*) multiplications, where *n* is the number of hospitals.

Once all the sketches are merged, the unary encoding has served its purpose. Then, we switch to a more standard ciphertext encoding of integers. To do that conversion, we just need to count the number of 0’s in the sketch vector, which we do by subtracting component-wise from a vector of all 1’s, and summing across the vector. We can thus sum up the total value in all merged buckets, all while remaining in ciphertext and only using a small number of additions and multiplications. At the end of the protocol, we decrypt the sum of all bucket values *N* and we estimate the cardinality using the regular LogLog estimation in plaintext [6]:

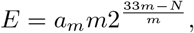

Where

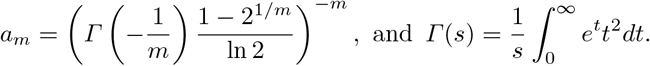

Since the number of buckets *m* is known and must be agreed upon to enact the protocol, the only sensitive information in that formula is *N*, the sum of all the bucket values. The information revealed by *E* is thus exactly equivalent to the information revealed by *N* . Hence, this entire computation reveals no information other than the answer to the query, and because it is precisely the LogLog algorithm shifted into ciphertext, the standard relative error is still 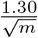.

### 2.2 Intersections through sampling

Private set intersection cardinality (PSI-CA), where parties want to jointly compute the intersection size between their sets without revealing anything else, has been extensively studied in the literature [3,4], including the multiparty variant [10]. However, unlike in prior work, we are not taking an intersection of sets belonging entirely to separate parties, but instead taking an intersection of sets that are themselves unions of the original data. Although it is possible to compute union sketches entirely in ciphertext, applying an intersection to them without decrypting is unfortunately not easy, even if we apply highly compressed intersection sketches [22], and we did not directly come up with a method of doing so.

Instead, we took an orthogonal approach based on sampling. Consider first taking an intersection of unions over a small population of total size *N*, where we have *n* hospitals collaborating. We can resolve this by simply using a bitvector of length *n*. Each hospital can then generate a bitvector *A*_*i,j*_ for hospital *i* and condition *j*, where *A*_*i,j*_[*k*] = 1 if and only if patient *k* has condition *j* at hospital *i*. Then we can create merged condition bitvectors *C*_*j*_ = *A*_1,*j*_ ∨ *A*_2,*j*_ ∨ … ∨ *A*_*n,j*_, where *C*_*j*_[*k*] = 1 if and only if patient *k* has condition *j* anywhere. The ∨ (OR) operator can be implemented by *X* ∨ *Y* = 1 − (1 − *X*) × (1 − *Y*) in homomorphic subtraction and multiplication. Afterwards, we can compute the intersection of the condition bitvectors by 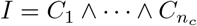, where the ∧ (AND) operator can be implemented by *A* ∧ *B* = *A* × *B*. Then, *I*[*k*] = 1 if and only if patient *k* has all of the conditions anywhere across the hospital system. After, we can do an internal sum of all set bits in the *I* vector to get the number of such patients. We can also easily generate a *U* vector by replacing the AND with an OR, to recover |*U* | the number of patients with any of the conditions.

Unfortunately, the universe of possible patients is too large for this to be currently feasible— because the hospitals do not want to reveal their patient populations, the bitvector would need length nearly 8 billion for the population of the planet (potentially less if geographically constrained, but still in the tens to hundreds of millions in many areas). If we could subsample, we might be able to use the bitvector technique on a smaller sample of patients, but we’d of course have to ensure that we sample the same sets of patients at each hospital. Another issue is that if we simply randomly sample patients, we might spend a large portion of our sample on patients that have nothing at all to do with the conditions we are interested in, especially if we are interested in looking at rare diseases.

Here, we make use of the union cardinality estimate *E* from the prior section, which we assume to be available in plaintext. Although no hospital knows the full list of patients that contributed to the union cardinality, notice that only patients who contributed to the union are potentially in the intersection. We can thus use *E* to compute a sampling rate *S/E*. Given a random hash function *h*_1_ (shared among the hospitals) mapping patient IDs to the unit interval [0, 1], we sample an ID if its hash function is less than the sampling rate. In expectation, we will then get a sample of *S* patients from the union, some fraction of which will be in the intersection. We then use some other random hash function *h*_2_ from patient IDs to [*m*]. For each condition *j*, hospital *i* creates a bitvector *A*_*i,j*_ of size *m*, and uses *h*_2_ to set bits for the appropriate sampled patients. That is, *A*_*i,j*_[*k*] = 1 if for some patient *x, h*_1_(*x*) *< S/E* and *h*_2_(*x*) = *k*. Afterwards, all of the *A*_*i,j*_ vectors can be merged as above, returning a count |*I*| of sampled patients in the intersection of all conditions and a count |*U* | of sampled patients in the union of all conditions. We then return 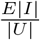 as the estimate of the total number of patients across the hospital network with all conditions.

#### Theorem 1

*The expected sample size S in a bitvector of length m should be chosen as S* = (*m*)^2*/*3^ *to minimize the error from hash collisions and sampling variance*.

*The expected standard error can then be overestimated at* 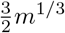.

*Proof*. There are two major sources of error in the returned count |*I*| of sampled patients in the intersection: (1) sampling variation and (2) errors due to hash collisions. Hash collisions are a problem because multiple patients may end up in the same bit within the bitvector, resulting in undercounting if both are in the intersection or overcounting if neither is in the intersection but between the two of them, they have all of the conditions. One solution would be to choose *m* = *O*(*S*^2^) large enough to avoid collisions entirely, but this is inefficient. Instead, we will strive to jointly minimize both types of errors.

First, note that the standard absolute error from sampling *S* items can be approximated by 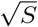. Second, we can upper bound the number of bins with multiple patients by simply the number of hash collisions. The expected number of hash collisions is 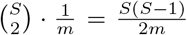. Thus, absolute error 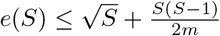, given fixed *m*.

Minimizing absolute error gives *S* = 0, which is not helpful, so instead we will minimize relative error 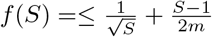 given fixed bitvector length *m*

Runtime of ciphertext operations

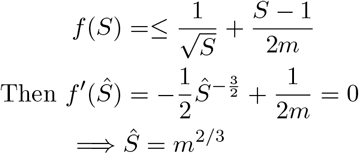

Furthermore, substituting this value back in gives an expected absolute error of

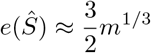

## 3 Results

### 3.1 Runtime benchmarks

We wrote benchmarking code (https://github.com/atleighton/rlwe-hll for union cardinality and https://github.com/yunwilliamyu/federated-approx-boolean for intersection sampling) to determine the single-threaded runtimes of the cryptographic steps within our protocol, using a 128-bit security parameter and varying the number of buckets and parties. We measure the total computation across all computing parties, referring to the central server as the “cloud” (Figure 3).

**Fig. 1.**
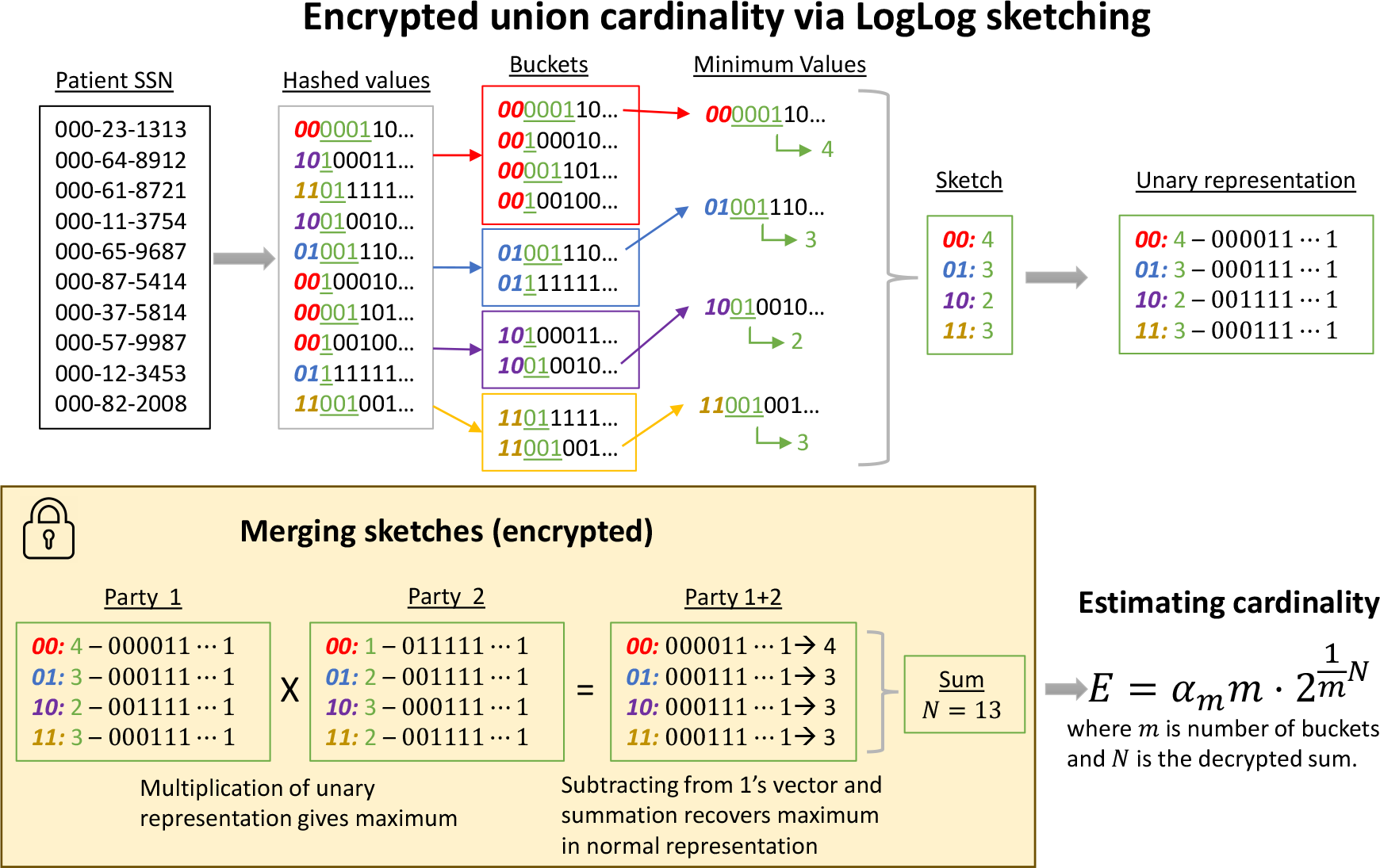
The encrypted union cardinality task can be solved via LogLog sketching and a unary encoding to allow for easy ciphertext maximums. Each party computes a ciphertext unary-encoded LogLog sketch, which is then sent an homomorphically merged to get just the raw value *N*, which can be used in conjunction with the number of buckets *m* to estimate union cardinality.

**Fig. 2.**
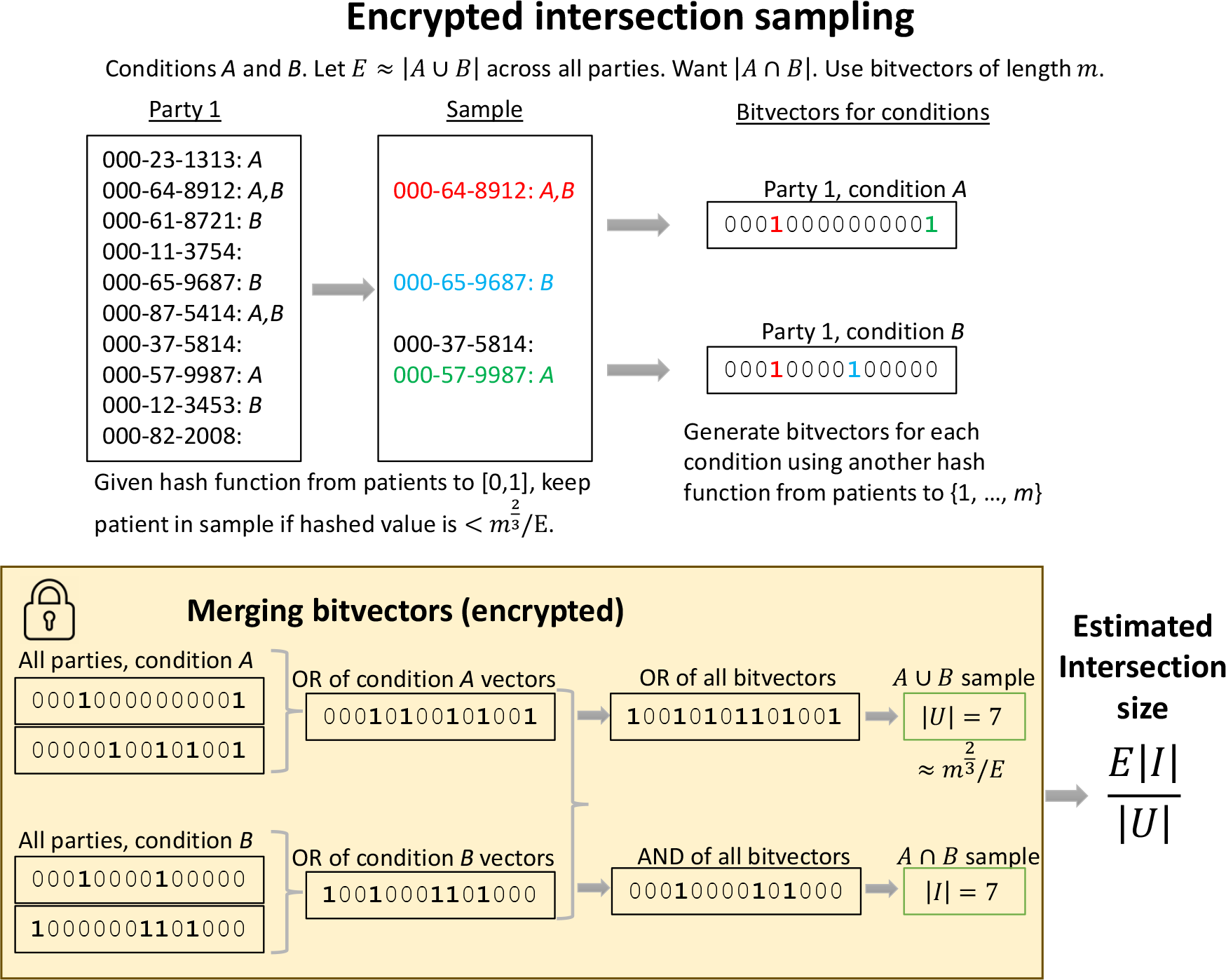
Given knowledge of an approximate size for the union of multiple conditions, we can use coordinated random sampling at a specified density to estimate the size of the intersection of those conditions. See Theorem 1 for the proof of a near-optimal sampling rate.

**Fig. 3.**
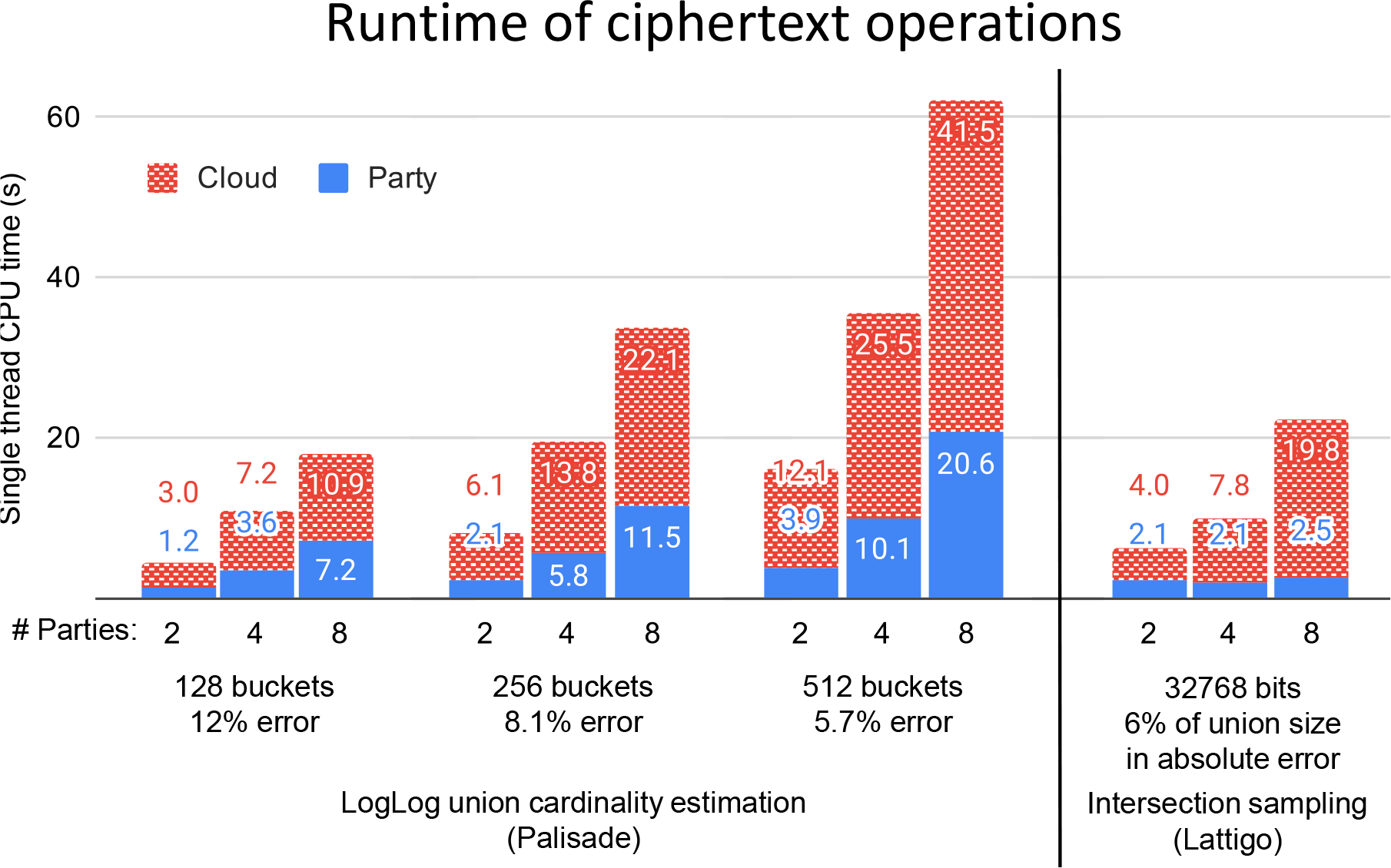
Single-thread runtime of ciphertext operations. (left) The LogLog union cardinality estimation implemented in Palisade increases linearly in runtime with number of buckets, and also number of buckets, for both the cloud and for all other parties (total). (right) The intersection sampling implemented in Lattigo empirically increases in runtime linearly to the number of parties; note that the party runtimes seems to be going up sublinearly as the number of parties increases, but we think this is simply a result of a large constant factor overhead in our implementation.

We implemented the LogLog union cardinality estimation in the Palisade C++ library. Even simulating up to 8 parties on a single CPU thread takes only just over one minute to return an estimate using 512 buckets (and commensurately less time for fewer buckets). With parallelization across cores on modern 32-core servers, ignoring network communication time, it is thus feasible to return queries in much less than a minute, on the order of seconds, making the method feasible for interactive query systems.

We also implemented intersection sampling using the Lattigo Golang library. To fully utilize the bit parallelism in the cryptosystem, we used a 32768 bit bitvector. For 8 parties on a single CPU-thread, this operation takes less than 25 seconds without parallelism. Per theorem 1, we downsample the density such that the union of all conditions should set roughly 1024 bits

### 3.2 Accuracy analysis

Our encrypted protocol for computing the union cardinality using a LogLog sketch exactly matches the unencrypted equivalent, and therefore has the same accuracy. The standard error of **LogLog** sketches is 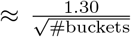, which is somewhat less efficient than e.g. the standard error of **Hyper-LogLog** approximations at 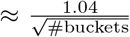 However, it follows simply we are able to achieve similar accuracy, regardless of our choice of sketching algorithms. In the **LogLog** case we increase the number of buckets by a factor of 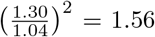. For the 512 bucket example, we end up with a 5.7% standard relative error in the union cardinality.

For the intersection sampling, we choose the sampling rate according to Theorem 1. Given a 2^15^ = 32768 bit vector, we therefore had in expectation 1024 bits set by the union of all conditions, and 48 incorrectly set bits, so ≤ 5% of the bits. This 5% error is of course scaled up by the density parameter through multiplying by the union cardinality estimate, so the total absolute final intersection error is roughly 5.985%≈ 6% of the union cardinality.

### 3.3 Security analysis

The security of our protocol in the semi-honest setting follows from the security of the well-known BFVrns homomorphic encryption scheme. Only three plaintext quantities are revealed: the bucket sum for the union cardinality LogLog sketch, the number of bits set from the subsampled union, and the number of bits set from the subsampled intersection. Together, these are used to compute the query outputs of the union and intersection cardinalities of the conditions that went into the query.

Importantly, in our protocol we do not seek to protect the outputs of the query from the adversary, only the underlying patient medical records. Thus, we are allowed to reveal the union and intersection cardinalities, as those are the answers to the queries. The union and intersection cardinalities correspond to the first and third quantities described above (after rescaling), so those quantities do not leak any information beyond the outputs of the queries. Also, the second quantity, the number of bits set from the subsampled union, will be a distribution centered around the expected sample size of *m*^2*/*3^. How far off the quantity is from the expected value may leak some small amount of information about which patients were subsampled by the hash function, but the set of potentially sampled patients is of course also fully revealed by the choice of hash function, which is public knowledge, so this leakage is minimal. Overall, this is just a restatement of the thesis of homomorphic encryption, which is that only the outputs of the algorithm are leaked in plaintext.

## 4 Discussion

A growing body of work aims to securely and efficiently compute statistics on medical data. Prior work has only addressed the federated union cardinality count query, but cannot be used for arbitrary Boolean queries including both ANDs and ORs (intersections and unions). We are the first to build a protocol capable of answering arbitrary Boolean queries across many computing parties, while having all the strong security guarantees of homomorphic encryption, revealing only three statistics, one of which is the query answer. Furthermore, our protocol is very fast, able to answer queries in a couple minutes without parallelism, and seconds with parallelism; these answers have error on the order of 6% of the union cardinality of all conditions considered, but that can of course be decreased with additional sampling in the standard fashion.

Beyond just prototyping a new system, our work demonstrates that modern homomorphic encryption libraries are sufficiently fast to be used in interactive federated queries, not just off-line analysis, resolving an open question about practically approximating federated Boolean count with almost no leakage of information. Furthermore, this manuscript shows that, provided that the correct algorithms and data representations are chosen, we can solve federated private Boolean count-queries that at first glance are not amenable to modern FHE based MPC frameworks.

Ultimately, we believe that the approach in this manuscript will become increasingly necessary as problems relating to the security and privacy of biomedical data grow in prevalence. Our work demonstrates that privacy trade-offs can be turned into computational power trade-offs. As computers become faster, we envision a future where patient privacy can be guaranteed even while aggregating their data for clinical research.

## 5 Acknowledgments

We thank Dr. Griffin Weber for introducing us to the federated count-query problem in addition to his guidance. We thank Dr. Isaac Kohane for his guidance and encouragement to think outside the box. Lastly, we thank Adam Sealfon for fruitful discussions.

